# The Cardiolipin Transacylase Tafazzin Regulates Basal Insulin Secretion and Mitochondrial Function in Pancreatic Islets from Mice

**DOI:** 10.1101/2021.01.15.426880

**Authors:** Laura K. Cole, Prasoon Agarwal, Christine Doucette, Mario Fonseca, Bo Xiang, Genevieve C. Sparagna, Nivedita Seshadri, Marilyne Vandel, Vernon W. Dolinsky, Grant M. Hatch

## Abstract

**Objective:** Tafazzin (TAZ) is a cardiolipin (CL) biosynthetic enzyme important for maintaining mitochondrial function. TAZ impacts both the species and content of CL in the inner mitochondrial membrane which are essential for normal cellular respiration. In pancreatic β-cells, mitochondrial function is closely associated with insulin secretion. However, the role of TAZ and CL in the secretion of insulin from pancreatic islets remains unknown.

**Methods:** Male 4-month-old doxycycline-inducible TAZ knock-down (TAZ KD) mice and wild-type littermate controls were utilized. Immunohistochemistry was used to assess β-cell morphology in whole pancreas sections, while *ex vivo* insulin secretion, CL content, RNA-Seq analysis and mitochondrial oxygen consumption were measured from isolated islet preparations.

**Results:** *Ex vivo* insulin secretion under non-stimulatory low-glucose concentrations was reduced ∼52% from islets isolated from TAZ KD mice. Mitochondrial oxygen consumption under low-glucose conditions was also reduced ∼58% in islets from TAZ KD animals. TAZ-deficiency in pancreatic islets was associated with significant alteration in CL molecular species and reduced oxidized CL content. In addition, RNA-Seq of isolated islets showed that TAZ KD increased expression of extracellular matrix genes which are linked to pancreatic fibrosis, activated stellate cells and impaired β-cell function.

**Conclusion:** These data indicate a novel role for TAZ in regulating normal β-cell function, particularly under low-glucose conditions.

## 1. Introduction

The global prevalence of type 2 diabetes (T2D) exacts an enormous burden on healthcare systems as well as amplifies morbidity and premature mortality [1]. One of the central components in the development and progression of T2D is the failure of pancreatic β-cells to secrete sufficient insulin [2]. In the presence of chronic fuel excess and obesity-associated insulin resistance, the β-cell responds with increased mass, insulin synthesis and secretory activity [2]. Diabetes develops when the compensatory measures cannot be sustained in response to the metabolic stress. While the specific molecular basis underlying the failure of pancreatic β-cells remains unclear, the intimate relationship between insulin secretion and mitochondrial function has been widely established [1].

Tafazzin (TAZ) is a transacylase which alters the content and molecular structure of cardiolipin (CL) in the inner mitochondrial membrane [3]. Specifically, TAZ promotes the enrichment of CL with linoleic acid yielding the dominant species tetralinoleoyl CL (L_4_CL). It is well established that CL content and molecular structure are required to maintain proper mitochondrial cristae architecture, and multiple biological processes including fatty acid metabolism, protein transport, apoptosis, and mitochondrial fission and fusion [3]. Moreover, the optimal activity of individual electron transport chain complexes as well as their assembly into respirasomes requires sufficient L_4_CL and total CL [3; 4]. The importance of TAZ and CL content in maintaining mitochondrial function is underscored by a rare disorder called Barth Syndrome in which patients lacking functional TAZ present with cardiomyopathy and skeletal dysfunction due to L_4_CL and total CL deficiency [5].

There are also reports that inadequate CL content may promote the development of mitochondrial dysfunction in pancreatic β-cells [6]. Mice lacking the phospholipid remodelling enzyme iPLA_2_β exhibit reduced insulin secretion from isolated islets as a result of increased oxidative damage to the mitochondrial membrane [6-8]. Based on the enrichment of CL in the mitochondria and sensitivity of polyunsaturated fatty acids (PUFA) to oxidative damage, the authors proposed that mitochondrial dysfunction could be a consequence of the inability to replace peroxidized fatty acid side chains [6; 7]. Consistent with the idea that iPLA_2_β deficiency increases β-cell dysfunction and failure, db/db mice express only low levels of iPLA_2_β in isolated pancreatic islets [8]. Nevertheless, the precise role of CL in maintaining β-cell function remains unclear since CL content nor degree of CL specific oxidation were determined in iPLA_2_β deficient islets. Furthermore, our previous work has indicated that the lack of TAZ is protective against the development of the metabolic syndrome in mice aged to 10-months of age [9]. In this report, we investigated whether CL remodelling is critical for maintaining β-cell mitochondrial function and insulin secretion in TAZ-deficient mice.

## 2. Materials and methods

### 2.1 Animal care

This study was performed with approval of the University of Manitoba Animal Policy and Welfare Committee which adheres to the principles for biomedical research involving animals developed by the Canadian Council on Animal Care and the Council for International Organizations of Medical Sciences. All animals were maintained in an environmentally controlled facility (22°C, 37% humidity, 12 h light/dark cycle) with free access to food and water. Experimental animals were generated by mating male transgenic (Tg) mice (B6.Cg-Gt(ROSA)26Sortm1(H1/tetO-RNAi:Taz,CAG-tetR)Bsf/ZkhuJ, Jackson Laboratory, Bar Harbour ME) which have a doxycycline (DOX) inducible TAZ-specific short-hair-pin RNA (shRNA) with female C57BL/6J mice (Jackson Laboratory, Bar Harbour, ME). The knock-down of TAZ was induced *in utero* by administering DOX (625 mg of DOX/kg of chow) as part of the standard low-fat 6% (w/w) fat rodent chow (Envigo Teklad, Rodent diet cat# TD.01306) to female C57BL/6 mice both before mating and during pregnancy. TAZ-deficiency is initiated *in utero* as previously described [10] to maximize the duration of TAZ-deficiency. The dams with litters continued to receive the DOX diet (TD.01306) to maintain TAZ KD in the offspring for the entire suckling period. Male offspring were weaned at 3 weeks of age onto the low-fat 6% (w/w) (TD.01306) DOX-containing diet (625 mg/kg diet). In addition, some dams and corresponding male offspring (Tg and Non-transgenic (NTg)) were fed a low-fat chow (6%, w/w) not treated with DOX which served to provide standard wild-type values. Male mice positive for the TAZ shRNA transgene were identified by PCR using primers (5’-CCATGGAATTCGAACGCTGACGTC-3’ forward, 5’-TATGGGCTATGAACTAATGACCC-3’ reverse) as described [9; 10].

### 2.2 Glucose tolerance test

Glucose tolerance tests (GTT) were performed as previously described [11]. The circulating concentration of insulin was determined from plasma separated from tail vein blood samples by centrifugation (10 min at 4000 rpm, 4°C) followed by a colorimetric ELISA assay (ALPCO, Salem NH, USA).

### 2.3 Pancreatic islet isolation and hormone assays

Mice were anesthetized with pentobarbital (110 mg/kg) and the pancreas perfused via the common bile duct with collagenase V as previously described [12]. Islets were manually picked using a dissecting microscope (SZ61, Olympus) and incubated overnight at 37°C, 5% CO_2_ in 10% FBS, 1% P/S RPMI 1640. The following day, islets from each animal were divided into 1.5 mL tubes (15 islets X 3 replicates). Insulin secretion experiments were performed by incubating each aliquot of islets at 37°C, 5% CO_2_ with Krebs-Ringer bicarbonate buffer (KRB) supplemented with 2.8 mM glucose for 1 h, followed by KRB containing 2.8 mM for 30 mins, 16.7 mM glucose for 30 mins, and KRB containing 16.7 mM glucose plus 3 mM KCL for 30 mins. The islets were then lysed (70% ethanol, 0.1 N HCl), dried down and resuspended in water. Total pancreatic islet insulin content, secreted insulin (ELISA, ALPCO), total pancreatic islet glucagon content and secreted glucagon (radioimmunoassay, Millipore) were quantitated and normalized to DNA content.

### 2.4 Histology and immunohistochemistry

The 4% paraformaldehyde (PFA)-fixed pancreas was cryopreserved, sectioned completely (12 µm) and mounted onto superfrost plus slides (Fisher). In order to assess islet morphology some sections were stained with cresyl violet. For single immunofluorescence (IF) analysis, sections were preincubated in blocking solution (PBS, goat serum 5% serum, 0.2% Triton X-100, 0.02% sodium azide, 0.1% BSA fraction V) for 2 hr at RT. Slides were then incubated with the corresponding primary IgG: rabbit glucagon (1:200), rabbit insulin (1:100), rabbit activated caspase 3 (1:200) (Cell Signaling Technologies, Danvers MA, USA) overnight at 4°C. Next day, after several washes in 1XPBS/Triton (PBS-T), slides were incubated in goat anti-rabbit Alexa-Fluor 488 or 594 conjugate (LifeTechnologies) in blocking serum buffer for 2 hr at RT. After several washes in PBS-T slides were then mounted with Vectashield/DAPI (Vector Laboratories). Signal for primary antibody 4-HNE (1:100) (Abcam) was amplified. Primary antibody for 4-HNE was incubated as described above. Biotinylated goat anti-rabbit IgG secondary antibody was added for 2 hr at RT. After PBS-T washes, streptavidin Alexa-Fluor conjugate 488 was added for an additional 2 hr at RT. Negative controls for insulin, glucagon, activated caspase-3 and 4-hydroxynonenal (4-HNE) staining were obtained in the absence of primary antibody. For assessment of fibrosis the pancreas was formalin-fixed and sectioned (5 µm) prior to trichrome or picro sirius red staining. Images were obtained using an Olympus IX70 inverted microscope and quantified using Image J software as previously described [12].

### 2.5 Mitochondrial analysis

Oxygen consumption rate (OCR) was measured using a Seahorse Bioscience Extracellular Flux Analyzer (model XF24, Agilent Technologies). Mouse islets (50 islets per well) were assayed in Seahorse XF media containing 2.8 mM glucose. Islets were then treated with 16.7 mM glucose followed by 10 µM oligomycin and finally 5 µM antimycin A + 5 µM rotenone. Basal oxygen consumption was considered to be the respiration sensitive to inhibition by 5 μM antimycin A + 5 μM rotenone. ATP-sensitive oxygen consumption was inhibited by 10 μM oligomycin, and ATP-insensitive respiration (heat) was the remaining proportion of basal oxygen consumption. Maximal oxygen consumption was achieved with 16.7 mM glucose. The addition of antimycin A inhibits all mitochondrial respiration therefore, any respiration in the presence of antimycin A was subtracted from all OCR values prior to calculating the average. The glucose-stimulated respiration was calculated by subtracting basal respiration from maximal respiration.

### 2.6 Quantitation of cardiolipin species

The molecular species of CL, oxidized CL (Ox-CL) and peroxidized CL (Per-CL) were quantitated from mouse islets using liquid chromatography coupled to electrospray ionization mass spectrometry (ESI-LCMS) in an API 4000 mass spectrometer (Sciex, Framingham, MA), as described previously [13]. Briefly, ∼300 islets per sample, pooled from ∼10 mice of the same genotype) lysed in PBS (200 µg protein) and lipids were extracted according to previously published methods [14; 15]. 1000 nmoles of tetramyristyl CL was used as an internal standard (Avanti Polar Lipids, Alabaster, AL, US). CL species were quantified per mg of protein based on a protein assay of homogenates (BCA, Pierce). Analysis was performed using the Analyst software with Ox-CL and Per-CL species determined by the M+16 and M+32 species respectively. While CL species had a retention time of approximately 8 min, Ox-CL and Per-CL had a retention time of 9-10 min using normal phase solvents. Ox-CL and Per-CL species underwent MS/MS to verify acyl composition. The total CL was calculated from the sum of the eight most prominent CL species (1442, 1424, 1448, 1450, 1472, 1474, 1496, 1498) which were previously defined by fatty acid side chain composition [16].

### 2.7 RNA-Seq library preparation and analysis

RNA was extracted from mouse islets (∼300 islets per sample, pooled from ∼10 mice of the same genotype) using RNAeasy Micro Kit (Qiagen, Valencia CA, USA) as per manufactures instructions. Ribosomal RNA was removed using Ribo-Zero rRNA removal kit (Illumina) and the purity and integrity assessed using Bioanalyzer 2000 (Agilent Technologies, Santa Clara CA, USA). The library was prepared using NEBNExt Ultra II Directional RNA Library Prep for Ilumina (NEB, Ipswich, MA, USA). Paired end RNA sequencing was performed using Illumina HiSeq 4000 PE100 in genome Quebec. The details of the sequencing reads are provided in Supplementary Table 1. The raw and processed data are submitted in GEO database with accession ID GSE136242. For the alignment of the reads default parameters of STAR version 020201 (PMID: 23104886) was used and the mouse reference genome version used was mm10. The differentially regulated genes were obtained by using HOMER v4.10 (PMID:20513432) script getDiffExpression.pl. The algorithm used for differentially regulated genes was R package DESeq2(PMID: 25516281).

### 2.8 RNA isolation and quantitative RT-PCR analysis

Total RNA was isolated from isolated pancreatic islets using a RNeasy® micro kit (Qiagen) and first-strand cDNA synthesis performed (0.5 µg total RNA) with SuperScript II (Invitrogen). PCR was performed using Eppendorf Realplex^2^ instrument with the gene specific primers as indicated (Supplementary Table 2). All mRNA levels were quantitated using a standard curve followed by normalization to *TfIIb*.

### 2.9 Statistical Analysis

Data are expressed as means ± standard error of the mean (SEM). Comparisons between animal treatment groups was performed by unpaired two-tailed Student’s t-test or analysis of variance using Tukey post-hoc analysis where appropriate. Student’s t-test was used for comparisons between low glucose and high glucose conditions within the same genotype. A probability value of <0.05 was considered significant. For RNA-Seq analysis, statistical analysis was performed using R packages DESEQ2 using the Bonferroni Hochberg method as previously described [12].

## 3. RESULTS

### 3.1 CL species are altered in pancreatic islets from TAZ deficient mice

The total CL content of isolated islets remained similar between the 4-month old TAZ KD (Tg+DOX) and wild-type animals (NTg+DOX, Fig. 1A). Mass spectrometry analysis indicated that several CL species were elevated in the TAZ KD mice including those (m/z 1472, 1474, 1496, 1498) which contain one or more very long chain PUFA (arachidonic acid, eicosapentaenoic acid) (Fig. 1 D). We confirmed that DOX feeding reduced TAZ in the pancreatic islets of TAZ shRNA Tg animals using the ratio of monolysocardiolipin (MLCL) to CL. Significant elevations in the MLCL/CL ratio are used to diagnose Barth syndrome patients due to the 100% sensitivity and specificity for the loss of adequate TAZ function [17]. We determined that the level of MLCL (359%) and the ratio of MLCL/CL (170%) were significantly elevated in the islets of TAZ KD mice compared to the NTg+DOX littermate controls (Fig. 1 B-C).

**Figure 1:**
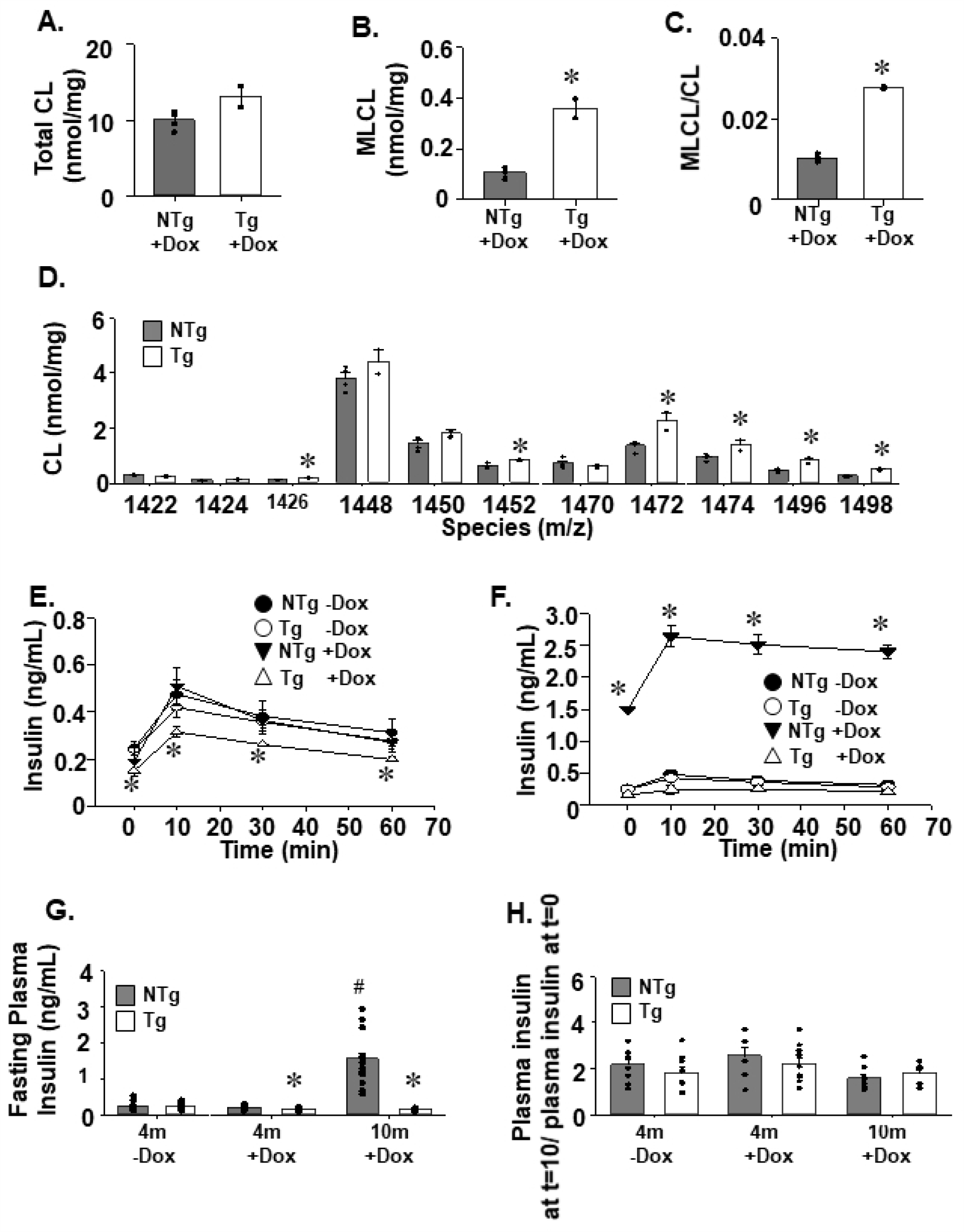
Tafazzin deficiency reduces plasma insulin levels. Islets isolated from Tg and NTg mice were used to quantitate total CL (A), MLCL (B) and the ratio of MLCL/CL (C) by mass spectrometry-mass spectrometry (n=2-4). Quantitation of individual CL species from isolated islets (D) (n=2-4). Plasma insulin levels following intraperitoneal glucose administration (2 g/mL) of Tg and NTg animals at 4 months (E) and 10 months (F) of age (n=8-10). Fasting plasma insulin levels (G). The plasma insulin level at t=10 minutes after glucose injection during GTT/plasma insulin level after 16 h fasting at t=0 minutes (before glucose injection) (H). Values are means ± SEMs. \**P*<0.05 compared with NTg mice fed the same diet. ^#^*P*<0.05 compared to all other groups.

### 3.2 TAZ deficiency reduces plasma insulin levels and *ex vivo* insulin secretion

Our previous work has indicated that TAZ-deficiency (Tg+DOX) is protective against the development of metabolic syndrome which occurred in aged littermate controls (10-month old NTg+DOX) [9]. Thus, it was expected that TAZ KD (Tg+DOX) mice may have reduced plasma insulin levels due in part to the development of hyperinsulinemia in control animals (10-month old NTg+DOX) which accompanies the elevated body weight and impaired glucose tolerance (Supplementary Fig. 1A, C) [9]. At 4-months of age the TAZ KD (Tg+DOX) mice had significantly reduced plasma insulin levels (∼25%) at all time points during GTT compared to littermate controls (NTg+DOX, Fig. 1E). This was complimented with small but significant increases (∼13%) of plasma glucose levels (Supplementary Fig. 1B). At 10-months, plasma insulin levels were also reduced during GTT in TAZ-deficient (Tg+DOX) mice compared to littermate controls (NTg +DOX, Fig. 1F). The downward shift in plasma insulin throughout the GTT for both 4 and 10-month-old TAZ KD mice was due to a significant reduction in baseline fasting plasma insulin levels (Fig. 1G) and not a reduction in the fold increase in insulin secretion stimulated by glucose (Fig. 1H).

In order to investigate whether the significantly lower plasma insulin levels observed in TAZ-deficient (Tg+DOX) mice were due to inherent low insulin secretion defect, we looked at insulin synthesis and secretion. We determined that the islet insulin content was maintained at a constant level when TAZ KD (Tg+DOX) mice were aged from 4 to 10 months (Fig. 2A). In contrast, insulin content increased (∼132%) in islets isolated from littermate controls as they aged from 4 to 10 months (NTg+DOX, Fig. 2 A). Both the % of insulin positive cells per islet and number of insulin positive cells per islet area were comparable between animal groups at both 4-month and 10-months of age (Fig. 2 B-D). Similarly, islet number and size were unaltered by TAZ-deficiency (Supplementary Fig. 2A-D). *Ex vivo* secretion of insulin under non-stimulatory low-glucose conditions (basal) was significantly reduced (∼48%) in the 4-month and 10-month TAZ KD mice compared to age matched wildtype controls (NTg+DOX, Fig. 2E and G respectively). However, there was no change in insulin secretion under high-glucose conditions or the insulin secretion index between genotypes for either age group (Fig. 2E-H). We also determined that the TAZ KD mice did not have dysfunctional insulin secretion machinery as membrane depolarization with KCL produced similar rates of insulin secretion between experimental groups (Supplementary Fig. 2E, F).

**Figure 2:**
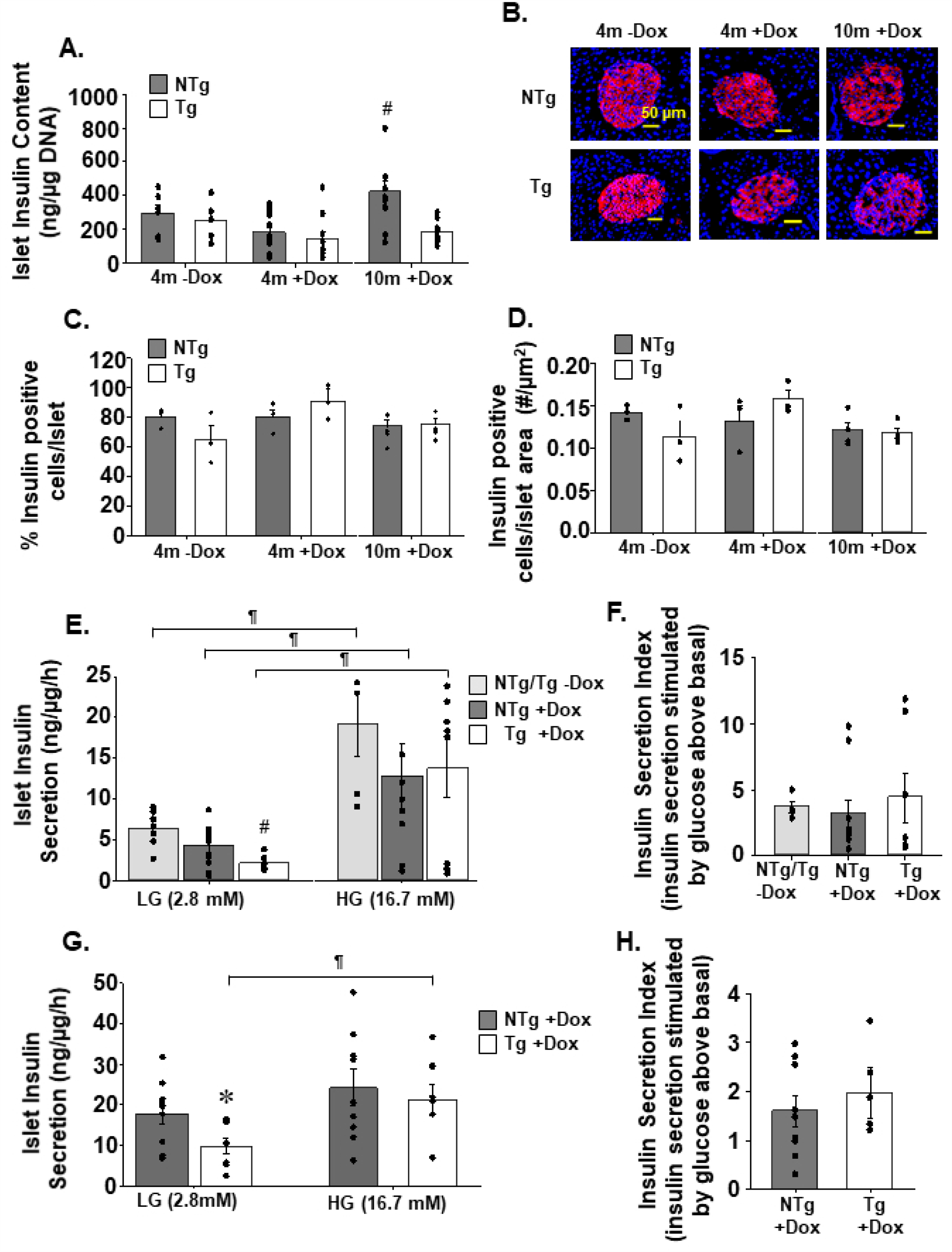
Mice lacking TAZ secrete less insulin during low glucose conditions. Islets isolated from Tg and NTg mice were used to quantitate total insulin content (n=5-11) (A). Representative immunofluorescent images (40X) of insulin-stained pancreas sections (B). The % of insulin positive cells per islet (C) and number of insulin positive cells per islet area (D) (n=3-4). Ex *vivo* insulin secretion by isolated islets from Tg and NTg mice at 4 months (E) and 10 months (G) of age in low-glucose (LG) and high-glucose (HG) conditions (n=5-10). The insulin secretion index (insulin secretion rate at high-glucose conditions /insulin secretion rate at low-glucose conditions) from islets isolated from 4 month (F) and 10 month (H) old mice. Values are means ± SEMs. \**P*<0.05 compared with NTg mice fed the same diet. ^#^*P*<0.05 compared to all other groups. ^¶^ *P*<0.05 compared with LG conditions from the same animal group.

### 3.3 Glucagon content is reduced but α-cells preserved in TAZ-deficient islets

Analysis of islet glucagon content revealed significant reduction in glucagon content in TAZ KD (Tg+DOX) animals at both 4 (∼56%) and 10-months (∼68%) of age compared to age matched wildtype controls (NTg+DOX, Fig. 3A). However, the reduction in glucagon content in TAZ KD mice did not correlate with reductions in either % glucagon positive cells per islet or number of α-cells per islet area (Fig. 3 B-D). Alternatively, TAZ-deficiency protected against the loss of α-cells which occurred in aged 10-month wildtype controls (Fig. 3B-D). *Ex vivo* secretion of glucagon from islets under low-glucose conditions (basal) and high-glucose conditions were similar between genotypes in both age groups (Fig. 3E, F). Fasting plasma glucagon levels were also constant between experimental groups (Fig. 3G).

**Figure 3:**
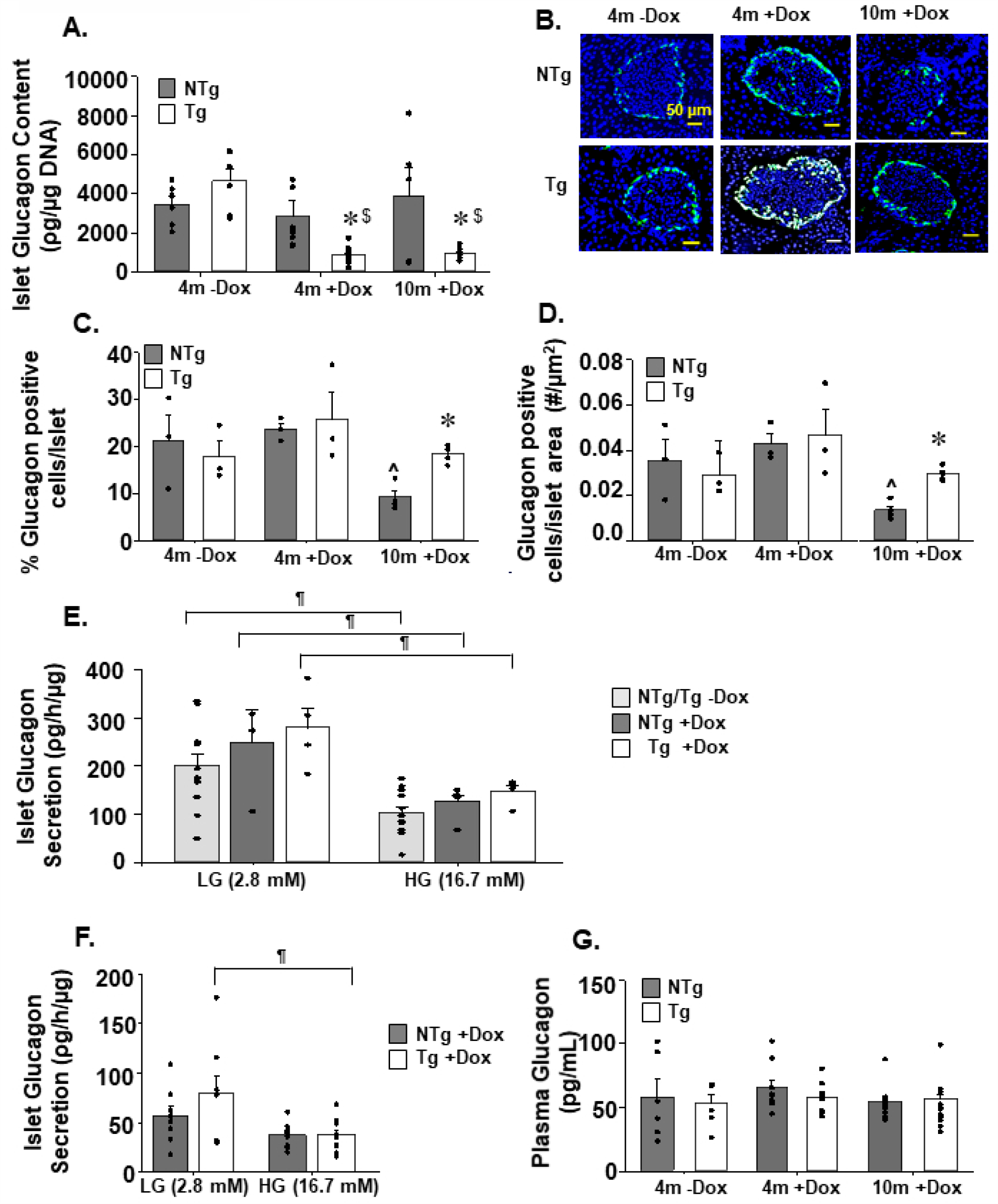
Analysis of pancreatic glucagon levels and secretion. Islets isolated from Tg and NTg mice were used to quantitate total glucagon content (A) (n=5-10). Representative immunofluorescent images (40X) of glucagon-stained pancreas sections (B). The % of glucagon positive cells per islet (C) and number of glucagon positive cells per islet area (D) (n=3-4). Ex *vivo* glucagon secretion by isolated islets from Tg and NTg mice at 4 months (E) and 10 months (F) of age in low-glucose (LG) and high-glucose conditions (HG) (n=4-12). (G) Fasting plasma glucagon levels (n=6-14). Values are means ± SEMs. \**P*<0.05 compared with NTg mice fed the same diet. ^$^*P*<0.05 compared to 4m-Dox of the same genotype. _^_*P*<0.05 compared to 4m+Dox of the same genotype. ^¶^*P*<0.05 compared with LG conditions from the same animal group.

### 3.4 TAZ-deficiency reduces basal respiration

We sought to determine whether the reduction of insulin secretion from islets isolated from TAZ KD (Tg+DOX) mice was associated with impaired mitochondrial respiratory function. Therefore, we measured OCR from islets in the presence of non-stimulatory low-glucose conditions. We determined that the level of basal OCR was significantly reduced (∼58%) in islets isolated from TAZ KD mice compared to the wildtype controls (NTg+DOX, Fig. 4A). This reduction was due to decreases in both the ATP-sensitive (∼83%) (oligomycin-sensitive) and ATP-insensitive (∼41%) oxygen consumption (heat) in the TAZ KD mice (Fig. 4B, C). Maximal respiratory capacity which was measured in the presence of high-glucose was comparable between TAZ KD mice and wildtype controls (NTg+DOX, Fig. 4D). Similarly, the glucose-stimulated respiration (Maximal OCR-basal OCR) was not altered by TAZ-deficiency (Fig. 4E). These data indicate that in TAZ KD mice, the attenuation of insulin secretion from islets incubated with low-glucose concentrations is associated with mitochondrial dysfunction.

**Figure 4:**
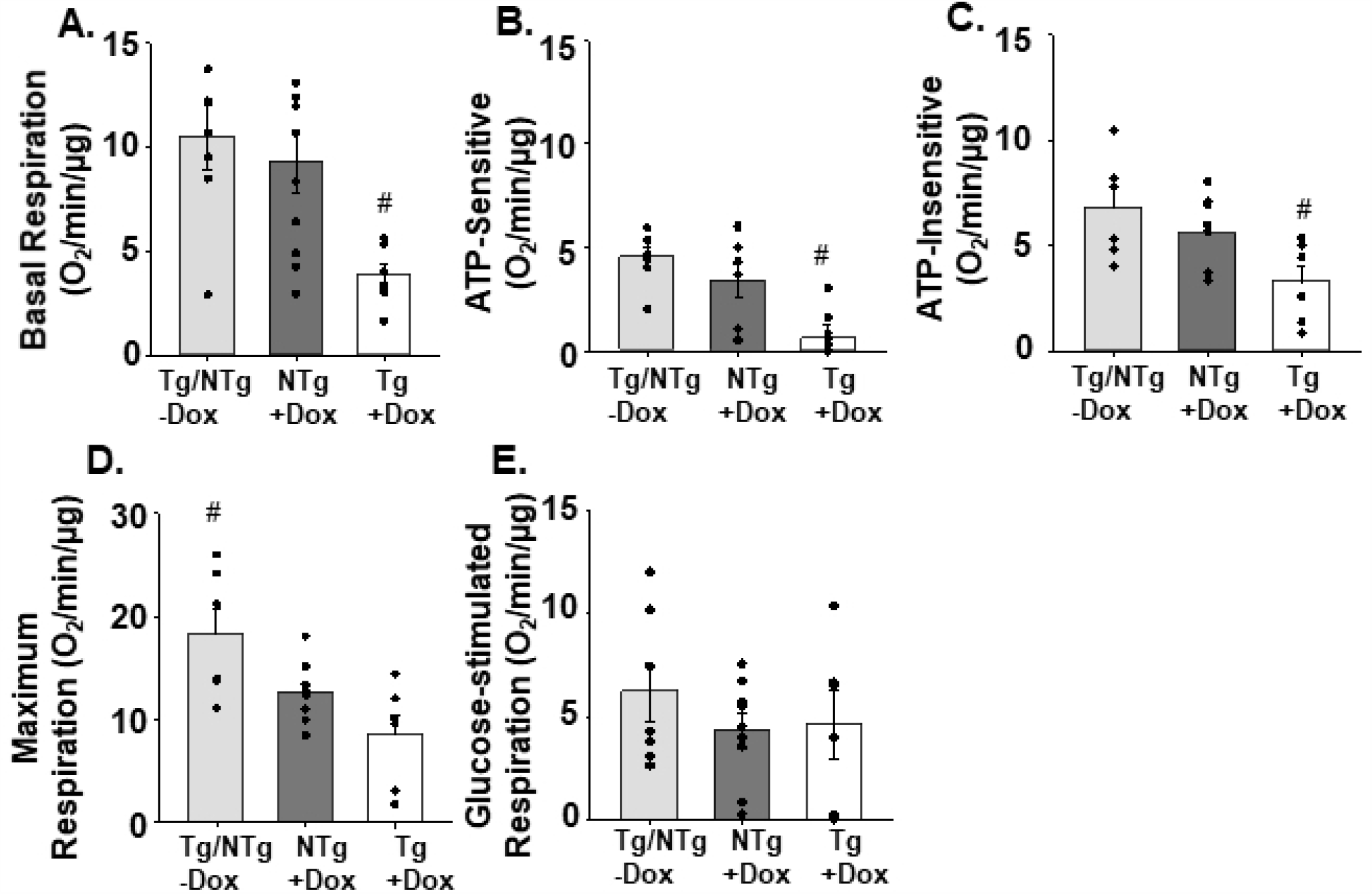
TAZ-deficiency reduced basal mitochondrial respiration in isolated islets. Basal mitochondrial respiration was measured from isolated islets in low-glucose (2.8 Mm, A). The ATP-sensitive (oligomycin-sensitive, B) and ATP-insensitive (i.e. heat, C) oxygen consumption rates. Maximum respiration in response to high-glucose (16.7 mM, D). Glucose-stimulated respiration ((glucose-stimulated) – (basal), E) (n=6-10). All measurements were normalized to µg of DNA. Values are means ± SEMs. ^#^*P*<0.05 compared to all other groups.

### 3.5 CL oxidation and peroxidation in TAZ-deficient islets

There are previous reports that lack of CL remodelling in pancreatic islets may promote mitochondrial dysfunction due to impaired replacement of peroxidized/oxidized fatty acid side chains [6; 7]. Therefore, we quantitated the level of oxidative damage to CL species using mass spectrometry. We determined that the amount of oxidized and peroxidized CL species were not different between 4-month-old TAZ KD (Tg+DOX) and wildtype control animals (NTg+DOX, Fig. 5A, C). Furthermore, significant decreases in oxidized (∼54%) (m/z 1472,1496, 1498) and peroxidized (∼38%) CL species (m/z 1496, 1498) were identified in TAZ-deficient islets when normalized to the corresponding unoxidized species (Fig. 5B, D).

**Figure 5:**
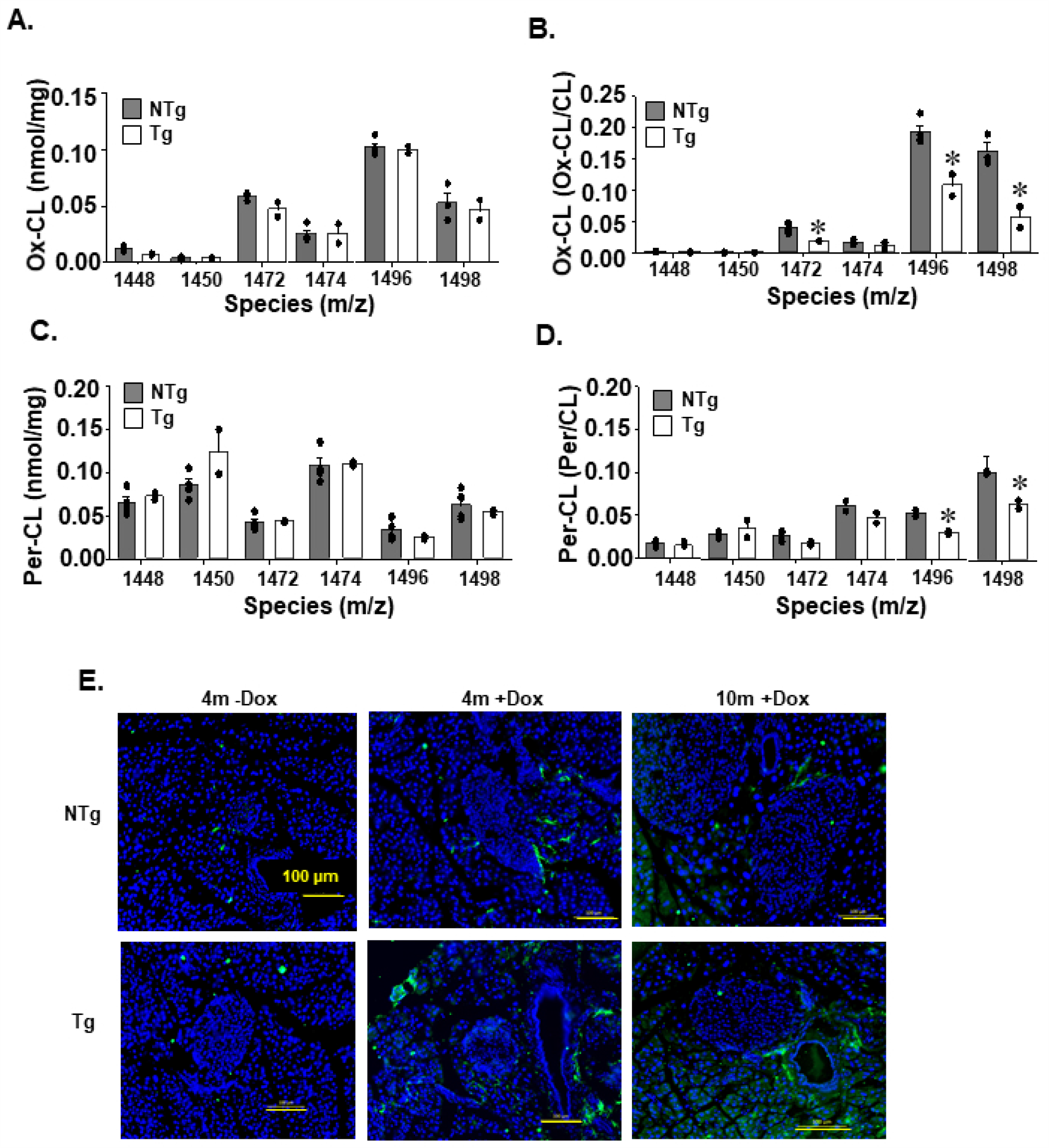
Quantitation of oxidized CL species in isolated pancreatic islets. Islets isolated from 4-month-old Tg and NTg mice were used to quantitate oxidized CL species by mass spectrometry-mass spectrometry (A) and expressed as a fraction of each individual CL species (B). Peroxidized CL species in islets were quantitated by mass spectrometry-mass spectrometry (C) and expressed as a fraction of each individual CL species (D). Values are means ± SEMs (n=2-4). Representative immunofluorescent (40X) images of 4-HNE stained pancreas sections (E) (n=3-4). \**P*<0.05 compared with NTg mice fed the same diet.

We also assessed the pancreatic level of 4-HNE, which is one of the dominant end-products of lipid peroxidation and mediators of oxidative stress-induced apoptosis [18]. Overall, minimal 4-HNE was detected within the islet and was similar across all experimental groups (Fig. 5E) (Supplementary Fig. 3). The amount of activated caspase-3, a final step in the apoptotic cascade, was also not different between islets from TAZ KD and wildtype littermate controls (Supplementary Fig. 4A, B). Together these data indicate that the mitochondrial dysfunction of islets from TAZ KD mice was not due to elevated lipid peroxidation, induction of apoptosis or an accumulation of oxidated CL species.

### 3.6 TAZ deficiency increases fibrosis in pancreatic islets

To elucidate the mechanism for altered insulin secretion and mitochondrial dysfunction in the TAZ KD mice we performed RNA seq on isolated pancreatic islets. Next-generation sequencing of mRNA from islets isolated from TAZ KD (Tg+DOX) and TAZ wildtype-controls (NTg+DOX) detected a total of 141 different genes (139 upregulated, 2 downregulated). Among the upregulated set of genes, *Fbln2, Fn1, Selp* and *Cdh6* showed the largest changes in expression (Fig. 6A, Supplementary Table 3). Novel pathways disrupted by TAZ KD were detected by functionally annotating differentially expressed genes using Ingenuity Pathway Analysis (Fig. 6B) The genes that were most highly represented clustered in the stellate cell activation and fibrosis pathways, which included various types of collagen and fibronectin (e.g., *Col1a1, Col5a1, Fn1*) (Supplementary Table 4). Enrichment of genes also occurred in the pathways associated with the biosynthesis of extracellular matrix components (epithelial mesenchymal transition pathway) and regulation of lipid metabolism (LXR/RXR activation) (Supplementary Tables 5&6).

**Figure 6:**
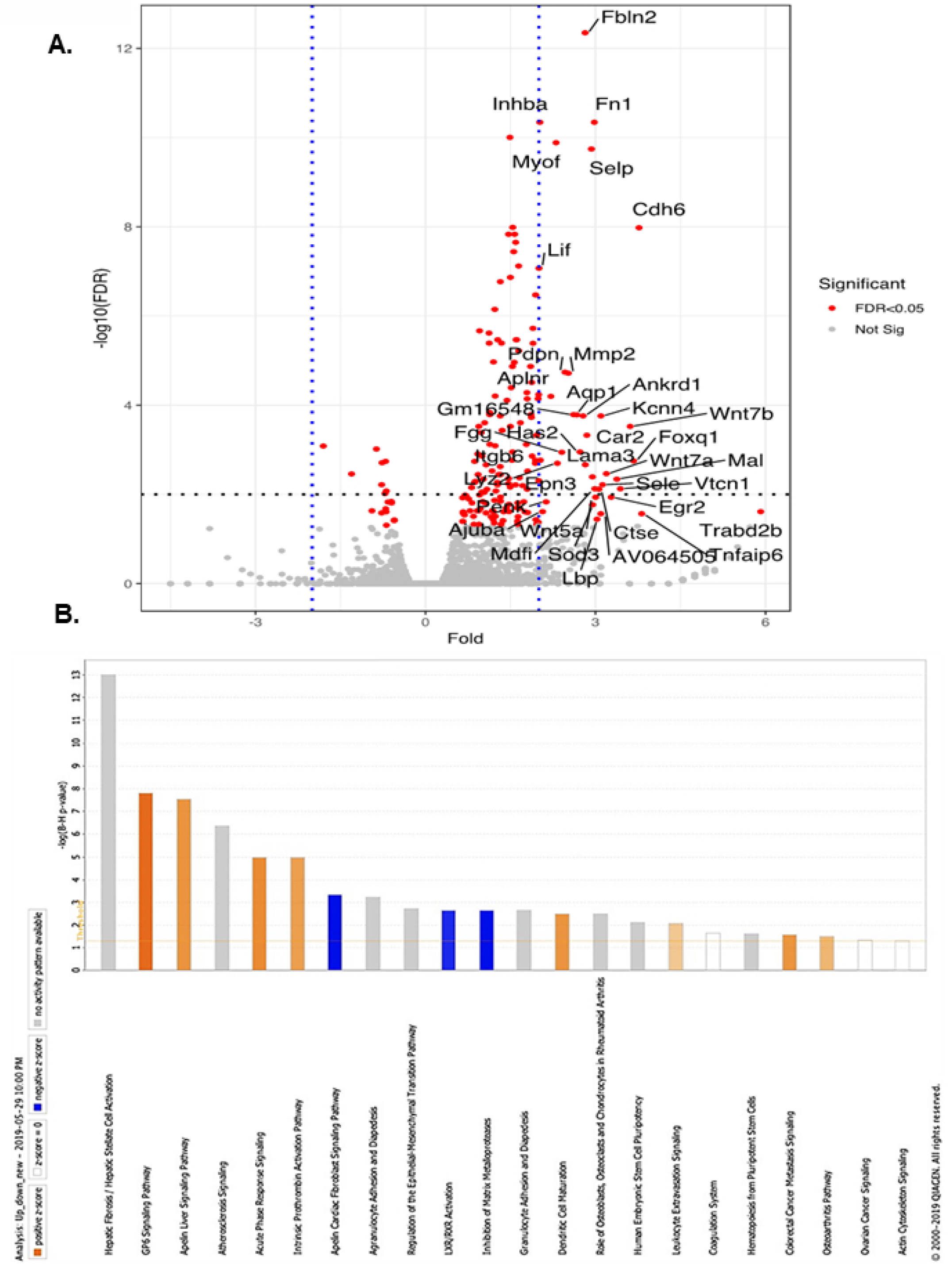
Transcriptomic profiling of TAZ KD islets revealed increased expression of extracellular matrix protein. Islet genes plotted as fold change against −log10(FDR) in a volcano plot. Significantly regulated genes (>1.5-fold) are marked in red (A). Highly enriched transcriptional pathways from the Ingenuity Pathway Analysis of mouse islet mRNA (B).

We performed a targeted assessment of TAZ and three genes with known roles in pancreatic fibrosis by qPCR analysis (*Fn1, Col6a2, Fbln2*) (Fig. 7B-D). We confirmed that *taz* gene expression levels was significantly reduced (∼89%) in islets isolated from TAZ KD mice (Fig. 7A). Several of the genes (*Fn1, Col6a2*) associated with fibrosis were differentially expressed with significant increases (2.8 to 10-fold) in the TAZ KD animals which is consistent with our RNA-Seq analysis (Fig. 7B-D). The extent of fibrosis occurring in the pancreas was further assessed by histological visualization (Fig. 7E). While in the controls picrosirius red and trichrome staining of collagen was limited to the blood vessels, islets derived form the TAZ KD mice displayed collagen along the periphery (Fig. 7E).

**Figure 7:**
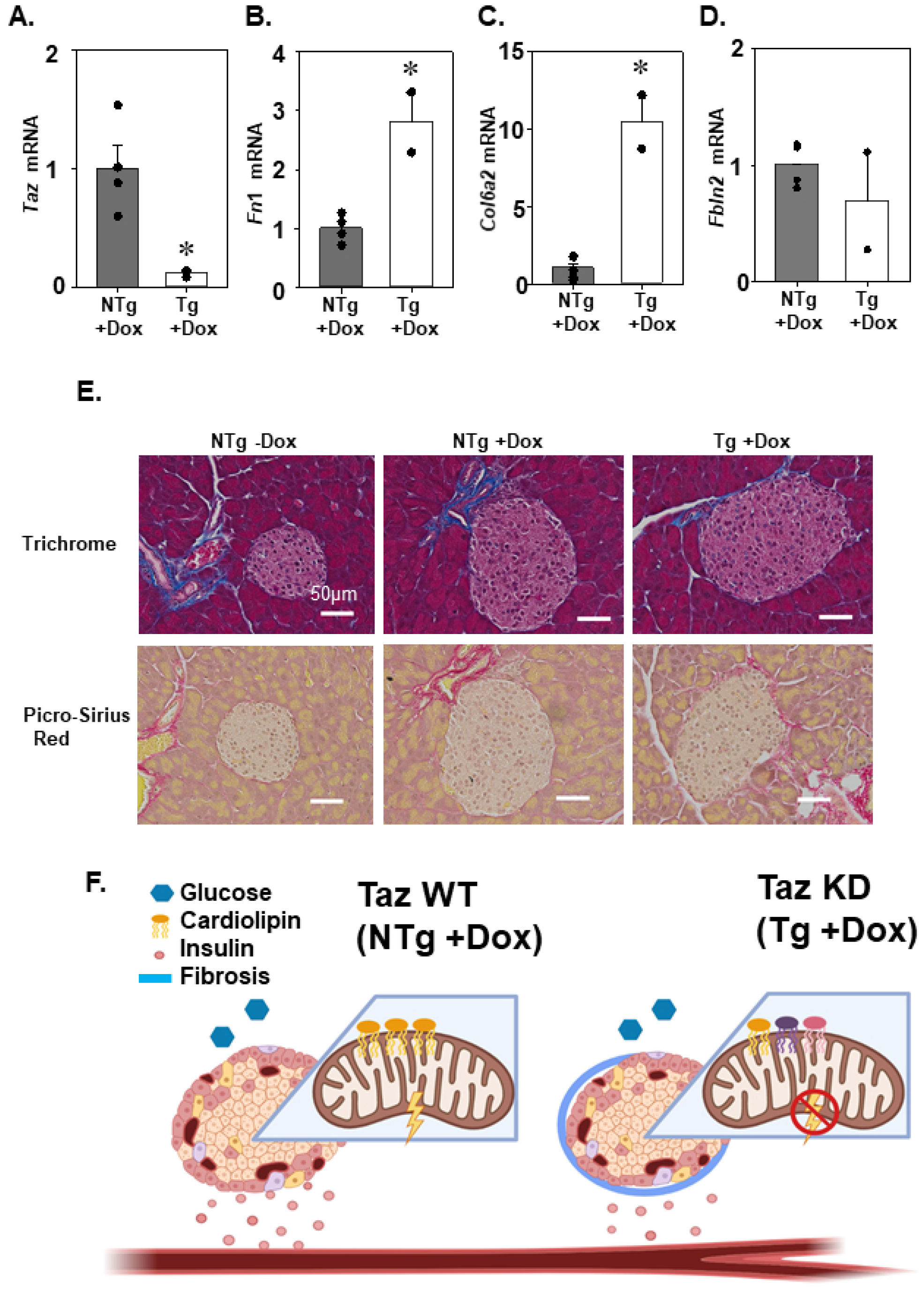
TAZ KD islets revealed increased expression of extracellular matrix protein. Quantitation of TAZ (*Taz*, A), fibronectin 1 (*Fn1*, B), collagen 6A2 (*Col6A2*, C) and fibulin 2 (*Fbln2*, D) mRNA by qPCR (n=2-4). Representative trichrome and picro-sirius red staining (40X) of pancreas sections (E) (n=4). Schematic summary: In NTg + Dox (TAZ WT) animals, low-glucose conditions are associated with a corresponding level of mitochondrial respiration and insulin secretion. In Tg + Dox (TAZ KD) animals, alteration in CL species (as indicated by different colors), increased fibrosis and reduced mitochondrial respiration are associated with a reduction of insulin secretion in low-glucose conditions (F). \**P*<0.05 compared with NTg mice.

## 4. DISCUSSION

The present study has defined a critical role for TAZ in regulating insulin release from the pancreatic β-cell. We demonstrated that TAZ-deficiency reduced *ex vivo* insulin secretion and mitochondrial oxygen consumption from isolated islets under non-stimulatory low-glucose concentrations. These effects were associated with significant increase in CL species containing very-long chain PUFA. Interestingly, our results also revealed that TAZ KD increased expression of ECM genes which are linked to pancreatic fibrosis, activated stellate cells and impaired β-cell function (Fig. 7F).

CL is almost exclusively localized to the inner mitochondrial membrane where it functions to maintain normal mitochondrial respiratory function. In the pancreatic islet mitochondrial oxidative phosphorylation is tightly linked to insulin release yet very little is known about the precise role of CL. Previously, it was demonstrated that inadequate CL remodeling in iPLA_2_β KO promotes pancreatic β-cell failure and reduces *ex vivo* insulin secretion [6-8]. We similarly determined that deficiency of the CL remodelling enzyme TAZ reduced insulin secretion from isolated pancreatic islets.

We demonstrated for the first time that a reduction in insulin secretion occurred in islets isolated from TAZ KD mice under basal conditions (low glucose levels). This result contrasts with the reduction in insulin secretion stimulated by high-glucose conditions identified in iPLA_2_β KO mice [7; 8]. One potential explanation for the difference between animal models is the divergence in underlying mechanisms. In the iPLA_2_β KO mice, the mechanism is characterized by oxidation of mitochondrial phospholipids and induction of apoptosis that reduced β-cell mass [8]. In addition, the inability to remove oxidized CL side chains has been proposed to increase production of 4-HNE and subsequently damage mitochondria in the iPLA_2_β KO mice [19]. Our data indicates that pancreatic islets isolated from the TAZ KD mice, exhibit normal β-cell mass, 4-HNE levels and caspase-3 activation as well as reduced levels of CL oxidation (Fig. 2 A-C, Fig. 5 A-E, Supplementary Fig 4A,B). Thus, indicating that a separate mechanism is responsible for the reduction in insulin secretion from islets isolated from TAZ KD animals.

Previously, we determined that TAZ-deficiency was protective against the development of obesity and hyperinsulinemia in aged 10-month-old animals [9]. Our current data supports a role for β-cell function in the protective effect of TAZ-deficiency. We have demonstrated for the first time that TAZ KD significantly reduces fasting plasma insulin levels due to reductions in β-cell insulin secretion under basal unstimulated conditions. Elevated basal insulin secretion rates are closely associated with hyperinsulinemia and the progression of T2D since they account for as much as half of the total daily insulin delivered [20].

Little is currently known about the mechanisms which control the rate of basal insulin secretion. Most processes identified focus on organization of the exocytotic machinery and regulation of ion channel activity. For example, there is evidence supporting the involvement of plasma membrane rafts, the organization of molecular scaffolding and gap junctions [21-24]. Alternatively, we have determined that the reduction in insulin secretion under low-glucose conditions correlated with a reduction in mitochondrial oxidative consumption rate. This is consistent with the connection between mitochondrial function and insulin secretion as well as the well-established role of TAZ in maintaining normal mitochondrial performance.

The prevailing model is that both CL content and acyl chain composition are critical for maintaining optimal mitochondrial function. Both reductions in CL content (>60%) and altered CL remodeling correlate with mitochondrial respiratory dysfunction in cardiac tissue from TAZ KD mice [25; 26], BTHS iPSC-derived cardiomyocytes [27; 28] and TAZ-deficient lymphocytes [29]. However, there is evidence that acyl chain remodeling may not be sufficient to regulate mitochondrial respiration. Mitochondrial respiratory dysfunction was not induced in yeast as a result of increased saturation of CL acyl chains [30], and in mice, high-fat diet-induced CL acyl chain remodelling was linked to diminished mitochondrial respiration in the liver, but not the heart [31; 32].

The specific acyl chain composition of CL has additionally been linked to the initiation of cellular effects through signaling functions [33]. For example, in response to acute injury, the initiation of apoptotic death signals was linked to specific CL oxidation products (linoleic, arachidonic acids and monolyso-CLs) across several tissues (brain, liver, heart, small intestine) [34]. Additional processes linked to CL signaling include mitophagy and immune responses (inflammasome activation, phagocytosis) although the specific acyl chains involved remain unknown [33]. Potentially, numerous cellular processes may be regulated by CL mediated signaling based on the vast number of species, oxidative metabolites and lipid mediators (>10^7^) [33]. However, there is a paucity of information on signaling pathways within the mitochondria, including those that regulate the post-translational modification of respiratory complexes [35]. Nevertheless, our results indicate that in mouse pancreatic islets, increased levels of very long PUFA-containing CL species were sufficient to impair oxidative phosphorylation in the presence of normal CL content. This finding underscores the sensitivity of pancreatic islets to CL molecular species and mitochondrial dysfunction.

Our data also indicates that islets isolated from TAZ-deficient animals may have elevated stimulation of stellate cells. Stellate cells have been extensively studied in the injured liver where they are the major ECM protein producing cell [36]. There is accumulating evidence for the role of stellate cells in the development of pancreatic fibrosis [37; 38] and in directly altering β-cell function [39]. Interestingly, pancreatic stellate cells were identified within the islets of healthy mice, particularly within the peripheral capsule [39] where fibrosis was localized was localized in the TAZ KD mice.

Islet fibrosis has been observed in patients [40] and rodent models of T2D [37; 41; 42]. Increased deposition of ECM components including collagen and fibronectin is consistent with the elevations observed in TAZ KD mice. However, the TAZ KD animals are actually protected against the development of metabolic syndrome [9]. This apparent contradiction may be explained due to the relatively small increase in fibrosis and ECM gene expression in the TAZ KD mice (∼1.1-3 fold) compared to animal diabetic models (2-30 fold) [37; 41; 42]. While the effect of ECM manipulations particularly within the peripheral capsule remains unclear [43], it is established that the relationship between the ECM and cellular function is dynamic. Therefore, the elevated ECM production in pancreatic islets from the TAZ KD mice may reflect the changes in cellular function (mitochondrial dysfunction) and/or promote changes in cellular function through surface receptors [44].

## 5. CONCLUSION

In conclusion, we have found that islets isolated from TAZ KD mice have reduced *ex vivo* insulin secretion and mitochondrial oxygen consumption under non-stimulatory low-glucose concentrations. Our findings also demonstrate that TAZ-deficiency in pancreatic islets was associated with significant alteration in CL molecular species, reduced oxidized CL content and increased expression of ECM genes. Together our data reveals a novel role for TAZ in maintaining normal β-cell mitochondrial function and insulin secretion. However, the protection offered by TAZ-deficiency simultaneously increases susceptibility to fibrosis which may have long-term consequences on insulin secretion.

## ACKNOWLEDGMENTS

PA is the recipient of a postdoctoral fellowship from Research Manitoba. LKC was the recipient of a CIHR/HSFC IMPACT Fellowship. GMH is a Canada Research Chair in Molecular Cardiolipin Metabolism. CAD is the Dr. J.A. Moorhouse Fellow of the Diabetes Foundation of Manitoba. VWD is the Allen Rouse-Manitoba Medical Services Foundation Basic Scientist. This research was supported by an Environments, Genes and Chronic Disease Canadian Institutes for Health Research (CIHR) Team Grant #144626, the Heart and Stroke Foundation of Canada, the Natural Sciences and Engineering Research Council (NSERC), Children’s Hospital Research Institute of Manitoba (CHRIM) and the University of Manitoba Research Grants Program (URGP). We thank McGill University and Genome Quebec Innovation Centre for performing the sequencing of the RNA libraries.

## ABBREVIATIONS

CL: Cardiolipin
ECM: Extracellular matrix
DOX: Doxycycline
HNE: Hydroxynonenal
GTT: Glucose tolerance test
NTg: non transgenic
OCR: Oxygen consumption rate
PLA: Phospholipase A
TAZ: Tafazzin
Tg: Transgenic
T2D: Type 2 diabetes

