## Supplementary Data for "The Cardiolipin Transacylase Tafazzin Regulates Basal Insulin Secretion and Mitochondrial Function in Pancreatic Islets from Mice"

**Supplementary Table 1** – Details of the RNA sequencing reads

| Sample | Total reads | Perfect aligned reads | Unmapped reads | Read pairs examined | Mapped pairs | Read pair duplicates | Percent duplication | Estimated library size |
|---|---|---|---|---|---|---|---|---|
| LC1 | 114,254,808 | 77,491,298 | 36,763,510 | 38,745,649 | 38,740,901 | 35,815,464 | 0.924 | 2,930,190 |
| LC2 | 129,323,672 | 91,163,760 | 38,159,912 | 45,581,880 | 45,577,263 | 42,727,046 | 0.937 | 2,854,834 |
| LC3 | 127,429,906 | 86,390,066 | 41,039,840 | 43,190,033 | 43,190,450 | 40,098,961 | 0.928 | 3,096,074 |
| LC4 | 127,369,080 | 88,221,550 | 39,147,530 | 44,106,038 | 44,106,038 | 40,979,417 | 0.929 | 3,131,360 |

**Supplementary Table 2** – Primers used for qPCR quantitation

| Gene | Primer name | Primer sequences | NCBI reference |
|---|---|---|---|
| *Fibronectin 1* | *Fn1 for* | gacccttacaggtttccca | NM_001276411 |
| | *Fn1 rev* | tgagcttaaagccagcgtca | |
| *Fibulin 2* | *Fbln2 for* | tagtcccgctcttctccaac | NM_001081437 |
| | *Fbln2 rev* | cctcctcgatgcagttctcc | |
| *Collagen 6a2* | *Col6a2 for* | gcccgtggacattgtcttcc | NM_146007 |
| | *Col6a2 rev* | aaggccacctgttgctgatt | |
| *Tafazzin* | *Taz for* | gaccctcatctctgggggat | NM_001173547 |
| | *Taz rev* | cagctccttggtgaagcaga | |
| *Transcription factor IIB* | *Tf11b for* | tggagatttgtccaccatga | NM_145546 |
| | *Tf11b rev* | gaattgccaaactcatcaaaact | |

**Supplementary Table 3 –** Top 20 increased gene expression in islets from TAZ KD mice compared to controls

| Gene symbol | Gene name | Fold-change | P-value |
|---|---|---|---|
| *Trabd2b* | trab domain containing protein 2B | 5.9 | 0.024 |
| *Tnfaip6* | tumor necrosis factor-inducible gene protein 6 | 3.8 | 0.027 |
| *Cdh6* | cadherin 6 | 3.8 | $1 \times 10^{-8}$ |
| *Foxq1* | forkhead box Q1 | 3.7 | 0.002 |
| *Wnt7b* | wingless 7b | 3.6 | 0.0003 |
| *Vtcn1* | V-set domain-containing T-cell activation inhibitor 1 | 3.4 | 0.007 |
| *Mal* | myelin and lymphocyte protein | 3.4 | 0.005 |
| *Egr2* | Early growth response protein 2 | 3.3 | 0.010 |
| *Wnt7a* | wingless 7a | 3.2 | 0.003 |
| *Sele* | E-selectin | 3.1 | 0.006 |
| *Kcnn4* | potassium/small conductance calcium activated channel, subfamily N, member 4 | 3.1 | 0.0002 |
| *AV064505* | non-coding RNA | 3.1 | 0.027 |
| *Ctse* | cathepsin E | 3.1 | 0.008 |
| *Lbp* | lipopolysaccharide binding protein | 3.0 | 0.036 |
| *Wnt5a* | wingless 5a | 3.0 | 0.012 |
| *Mdfi* | myoD family inhibitor | 3.0 | 0.007 |
| *Fn1* | fibronectin 1 | 3.0 | $5 \times 10^{-11}$ |
| *Sod3* | superoxide dismutase 3 | 3.0 | 0.017 |
| *Epn3* | epsin 3 | 2.9 | 0.004 |
| *Selp* | selectin P | 2.9 | $2 \times 10^{-10}$ |
| *Car2* | carbonic anhydrase 2 | 2.8 | 0.0005 |

**Supplementary Table 4**- Enrichment of fibrosis/hepatic stellate cell activation genes in islets from TAZ KD mice.

| Gene symbol | Gene name | Fold-change | P-value |
|---|---|---|---|
| *Col1a1* | collagen type 1, alpha 1 | 1.5 | $1 \times 10^{-10}$ |
| *Col1a2* | collagen type 1, alpha 2 | 1.6 | $3.4 \times 10^{-6}$ |
| *Col3a1* | collagen type 3, alpha 1 | 1.6 | $2.2 \times 10^{-8}$ |
| *Col5a1* | collagen type 5, alpha 1 | 1.2 | 0.036 |
| *Col5a2* | collagen type 5, alpha 2 | 1.3 | 0.0002 |
| *Col5a3* | collagen type 5, alpha 3 | 1.8 | $7.2 \times 10^{-5}$ |
| *Col6a2* | collagen type 6, alpha 2 | 1.8 | 0.003 |
| *Col15a1* | collagen type 15, alpha 1 | 1.6 | $3.7 \times 10^{-8}$ |
| *Col18a1* | collagen type 18, alpha 1 | 1.4 | $4.1 \times 10^{-6}$ |
| *Ctgf* | connective tissue growth factor | 1.6 | $1.1 \times 10^{-5}$ |
| *Fn1* | fibronectin 1 | 3.0 | $5 \times 10^{-11}$ |
| *Il6* | interleukin 6 | 1.9 | 0.002 |
| *Lbp* | lipopolysaccharide binding protein | 3.0 | 0.036 |
| *Mmp2* | matrix metalloproteinase 2 | 2.5 | $2 \times 10^{-5}$ |
| *Pdgfb* | platelet-derived growth factor subunit B | 1.1 | 0.0001 |
| *Pdgfrb* | platelet-derived growth factor receptor beta | 1.1 | 0.008 |
| *Serpin E1* | plasminogen activator inhibitor 1 | 1.2 | $1.1 \times 10^{-5}$ |
| *Timp1* | tissue inhibitor of metalloproteinase 1 | 1.8 | 0.0008 |

**Supplementary Table 5**- Enrichment of epithelial-mesenchymal transition pathway genes in islets from TAZ KD mice.

| Gene symbol | Gene name | Fold-change | P-value |
|---|---|---|---|
| *Jag1* | jagged 1 | 1.0 | 0.005 |
| *Mmp2* | matrix metalloproteinase 2 | 2.5 | $2 \times 10^{-5}$ |
| *Notch2* | notch 2 | 1.2 | 0.03 |
| *Pdgfrb* | platelet-derived growth factor receptor beta | 1.1 | 0.008 |
| *Wnt5a* | wingless 5a | 3.0 | 0.012 |
| *Wnt7a* | wingless 7a | 3.2 | 0.003 |
| *Wnt7b* | wingless 7b | 3.6 | 0.0003 |
| *Zeb2* | zinc finger E-box binding homeobox 2 | 1.6 | 0.01 |

**Supplementary Table 6**- Enrichment of LXR/RXR activation genes in islets from TAZ KD mice.

| Gene symbol | Gene name | Fold-change | P-value |
|---|---|---|---|
| *C3* | Complement component 3 | 1.2 | 0.02 |
| *Il6* | interleukin 6 | 1.9 | 0.002 |
| *Lbp* | lipopolysaccharide binding protein | 3.0 | 0.036 |
| *Lyz2* | lysozyme | 2.3 | 0.002 |
| *Ptgs2* | prostaglandin endoperoxide synthase 2 | 1.5 | 0.0003 |
| *Serpin F1* | pigment epithelium-derived factor | 1.8 | 0.025 |

Supplementary Figure 1

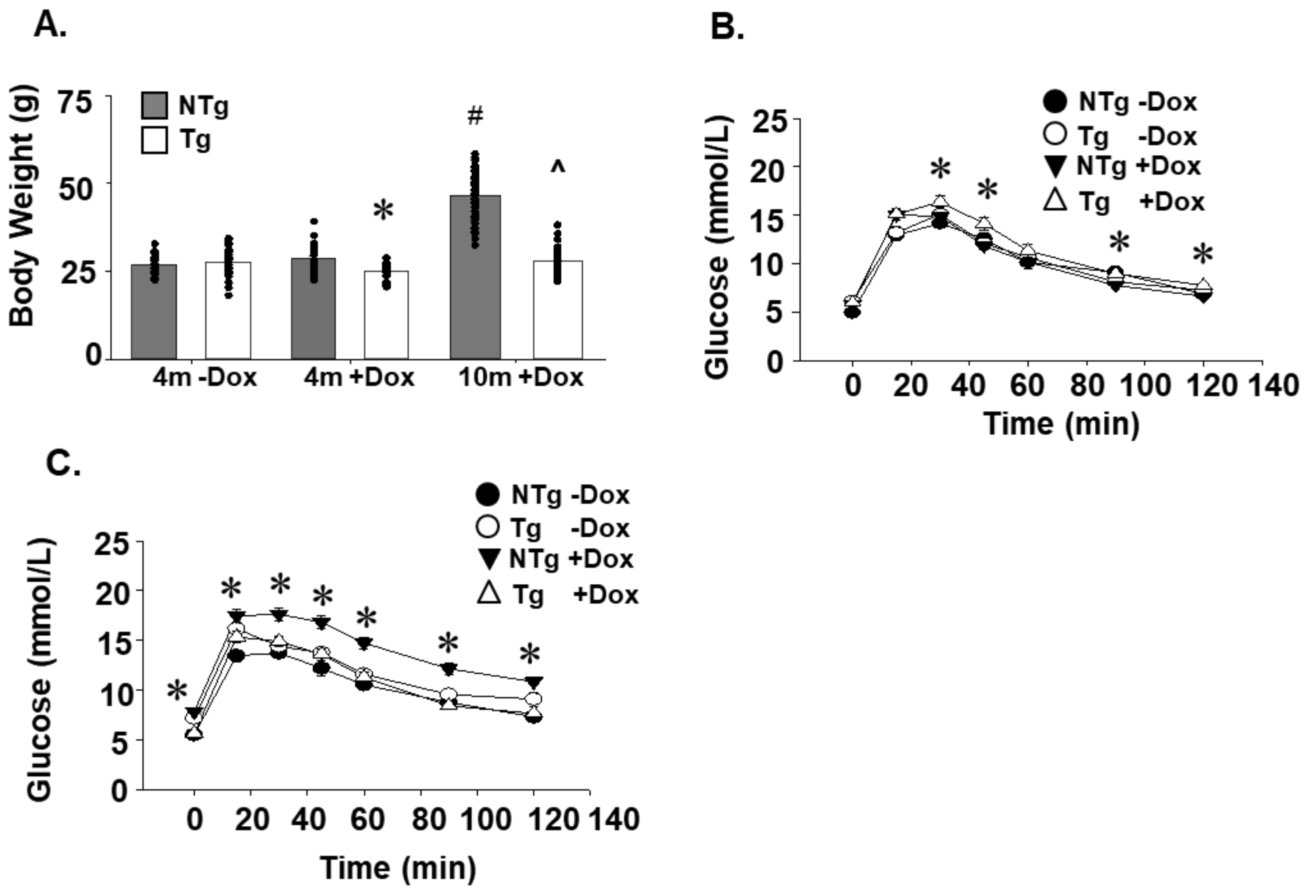


Supplementary Figure 1: Body weights and analysis of glucose tolerance in TAZ deficient mice. The body weights of NTg and Tg mice (A) (n=24-70). Glucose tolerance tests performed in NTg and Tg mice at 4 months (B) and 10 months (C) of age (n=12-28). Values are means ± SEMs. *$P$<0.05 compared with NTg mice fed the same diet. ^$P$<0.05 compared to 4m+DOX of the same genotype. $^{\#}P$<0.05 compared to all other groups.

Supplementary Figure 2

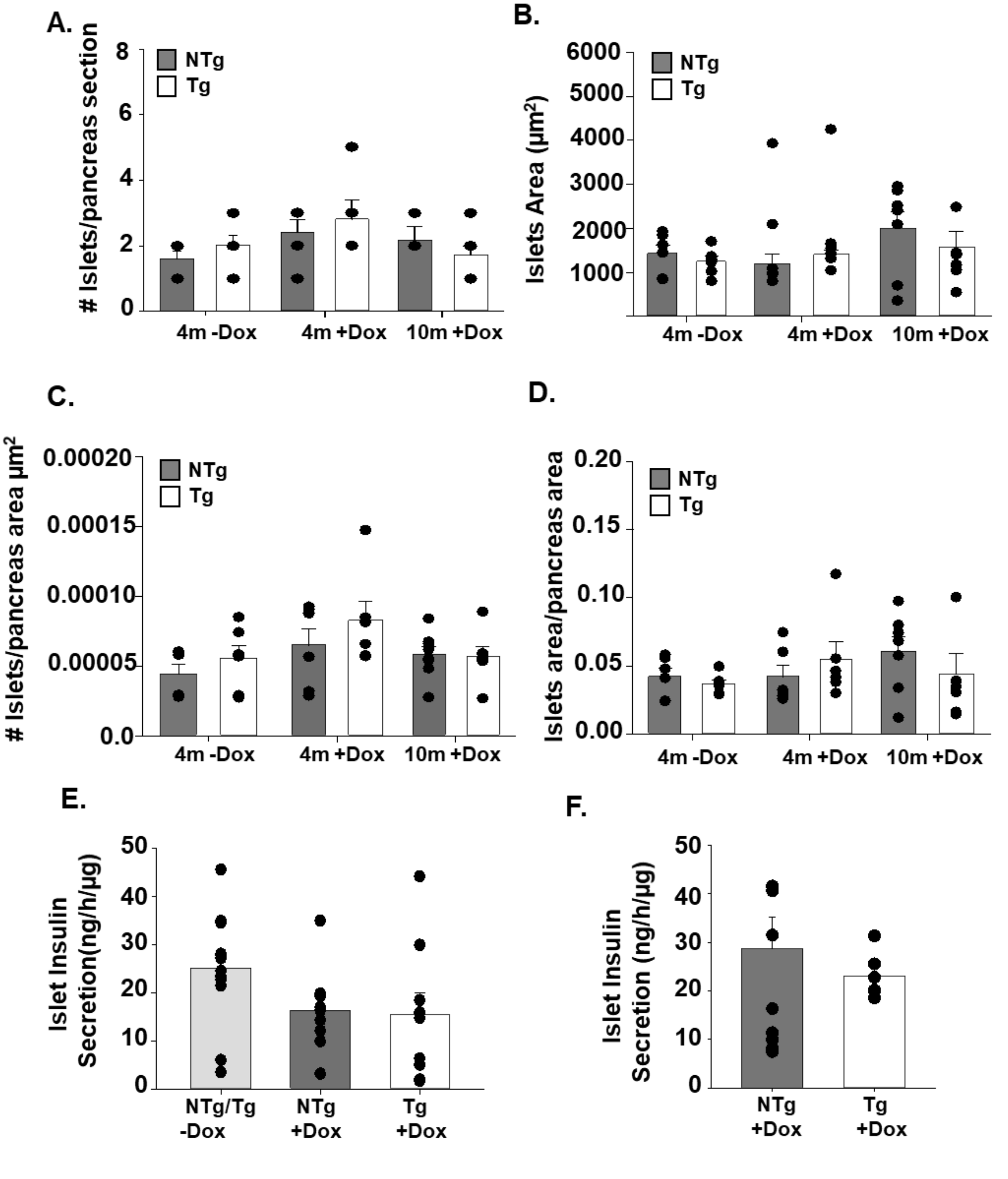


Supplementary Figure 2: Assessment of pancreas morphology in TAZ deficient mice. Number of islets per pancreas section (A). Islet area (B). Number of islets per pancreas tissue area (C). Islet area per pancreas tissue area (D). Ex *vivo* insulin secretion by isolated islets from Tg and NTg mice at 4-months (E) and 10-months (F) of age in high-glucose (16.7 mM) + 3 mM KCL conditions (n=5-10). All values are means ± SEM (n=6-7). Values are means ± SEMs.

Supplementary Figure 3

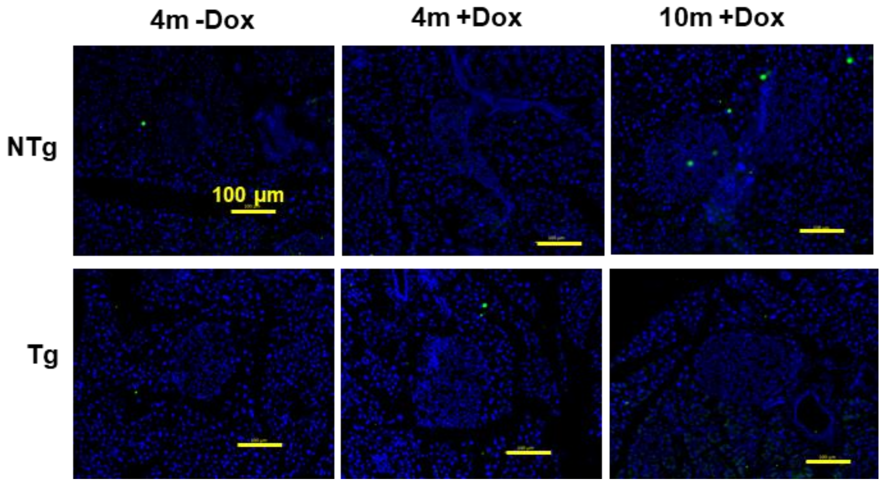


Supplementary Figure 3: Negative controls for 4-HNE staining of pancreas sections. Representative immunofluorescent images (20X) of 4-HNE stained pancreas sections in the absence of primary antibody (n=3-4).

### Supplementary Figure 4

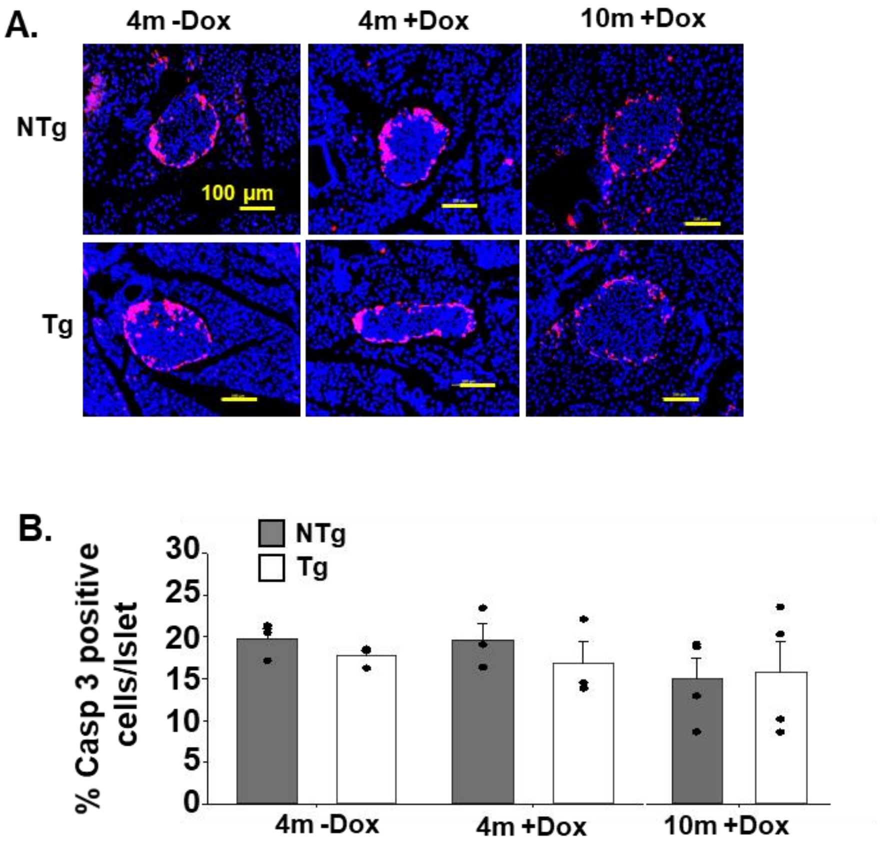


Supplementary Figure 4: TAZ deficiency does not increase activation of caspase-3 in islets. Representative immunofluorescent images (20X) of activated caspase-3 stained pancreas sections (A) and corresponding quantitation of the % caspase-3 positive cells per islet (B). All values are means ± SEM (n=3-4).
